# Early-life stress lastingly impacts microglial transcriptome and function under basal and immune-challenged conditions

**DOI:** 10.1101/2022.07.13.499949

**Authors:** Kitty Reemst, Laura Kracht, Janssen M. Kotah, Reza Rahimian, Astrid A.S. van Irsen, Gonzalo Congains Sotomayor, Laura Verboon, Nieske Brouwer, Sophie Simard, Gustavo Turecki, Naguib Mechawar, Susanne M. Kooistra, Bart J. L. Eggen, Aniko Korosi

## Abstract

Early-life stress (ELS) leads to increased vulnerability to psychiatric disorders including depression later in life. Neuroinflammatory processes have been implicated in ELS-induced negative health outcomes, but how ELS impacts microglia, the main tissue-resident macrophages of the central nervous system, is unknown. Here, we determined the effects of ELS induced by limited bedding and nesting material during the first week of life (postnatal days [P]2 – 9) on microglial i) morphology; ii) hippocampal gene expression; and iii) synaptosome phagocytic capacity in male pups (P9) and adult (P200) mice. The hippocampus of ELS-exposed adult mice displayed altered proportions of morphological subtypes of microglia, as well as microglial transcriptomic changes related to the tumor necrosis factor response and protein ubiquitination. ELS exposure leads to distinct gene expression profiles during microglial development from P9 to P200 and in response to an LPS challenge at P200. Functionally, synaptosomes from ELS-exposed mice were phagocytosed less by age-matched microglia. At P200, but not P9, ELS microglia showed reduced synaptosome phagocytic capacity when compared to CTRL microglia. Lastly, we confirmed the ELS-induced increased expression of the phagocytosis-related gene *GAS6* that we observed in mice, in the dentate gyrus of individuals with a history of child abuse using *in situ* hybridization. These findings reveal persistent effects of ELS on microglial function and suggest that altered microglial phagocytic capacity is a key contributor to ELS-induced phenotypes.

## Introduction

Exposure to early-life stress (ELS) has long lasting effects on brain structure and function and increases the risk for psychiatric illness later in life (1–3). Human and rodent studies have demonstrated that stress during sensitive developmental periods impacts mood (2–5), cognition (6–10) and the neuroimmune system(11–16).

While the mechanisms for this early-life programming of later-life health remain poorly understood, there is increasing evidence that long-term impact on the neuroimmune systems and microglia in particular might contribute to these effects (17,18). Microglia are innate immune cells in the brain parenchyma that can respond to environmental cues such as stress by means of cytokine release and phagocytosis (19–22) and are crucial for proper brain development and function by e.g. synaptic pruning (23–25).

There is ample evidence from maternal inflammation and early-life infection studies in rodents that early experiences can enduringly change microglial phenotypes (26). This is thought to be mediated via epigenetic mechanisms that reinforce microglial training or desensitization, i.e. hyper- or hyposensitivity, towards secondary inflammatory challenges in later life (15,27–32). Considering the well-documented interactions between stress and inflammation (33–35), ELS might, similarly, program microglia (36,37). In fact, we and others have previously shown age-dependent effects of ELS on microglia that are largely based on morphological characterization at basal (14,38,39) and challenged states, e.g. in response to amyloid-β pathology (14). While transcriptomic studies of microglia in the context of ELS are rare, microglial gene expression profiling shortly after postnatal stress revealed ELS alterations chemotactic and phagocytic processes (13). However, a thorough understanding of ELS’ short and long-term impact on microglial gene expression and function, and whether such changes also occur in the human brain is currently lacking. Therefore, we studied 1) the immediate effects of ELS on microglial gene expression; 2) the long-term effects of ELS on microglial morphology and gene expression profile in mice, in basal and immune-challenged states, in order to unmask potentially latent impacts of ELS; 3) the implications of these alterations for microglial phagocytic capacity; and 4) whether one of our target genes is similarly altered in the human condition.

We here demonstrate for the first time that ELS leads to long-term changes in the microglial transcriptome at P200, modifies the trajectory of gene expression changes during development and in response to LPS, and reduces microglial phagocytosis of synapses at P9 and P200. Finally, we validate in a postmortem human cohort that *GAS6*, a phagocytosis-related gene found upregulated in ELS mice, is also increased in the hippocampal microglia of individuals with a history of child abuse.

## Materials & Methods

### Experimental design, breeding, and early-life stress model

To determine acute and long-term effects of ELS on microglia, we exposed seven different cohorts of mice to ELS or control (CTR) conditions from postnatal days (P)2 to P9 via the limited bedding and nesting model (see Supplementary Methods). We confirmed across the generated cohorts for this study the reduction in body weight gain between P2 and P9 in ES-exposed mice (Figure S1A) (38,40).

Cohort 1 was used for morphological characterization of microglia 24 hours after PBS or LPS (5 mg/kg) injection at age of 3-5 months (*CTR-PBS*: n=5, *ELS-PBS*: n=4, *CTR-LPS*: n=10, *ELS-LPS*: n=6). Cohort 2 was sacrificed at P9, while cohort 3 was allowed to mature until P200 and injected with either PBS or LPS (1 mg/kg). Microglia from cohorts 2 and 3 were isolated for transcriptomic characterization (*P9-CTR:* n=12, *P9-ELS:* n=12, *P200-CTR-PBS:* n=7, *P200-ELS-PBS:* n=6, *P200-CTR-LPS*: n=7, *P200-ELS-LPS*: n=5, Figure 1A).

**Figure 1:**
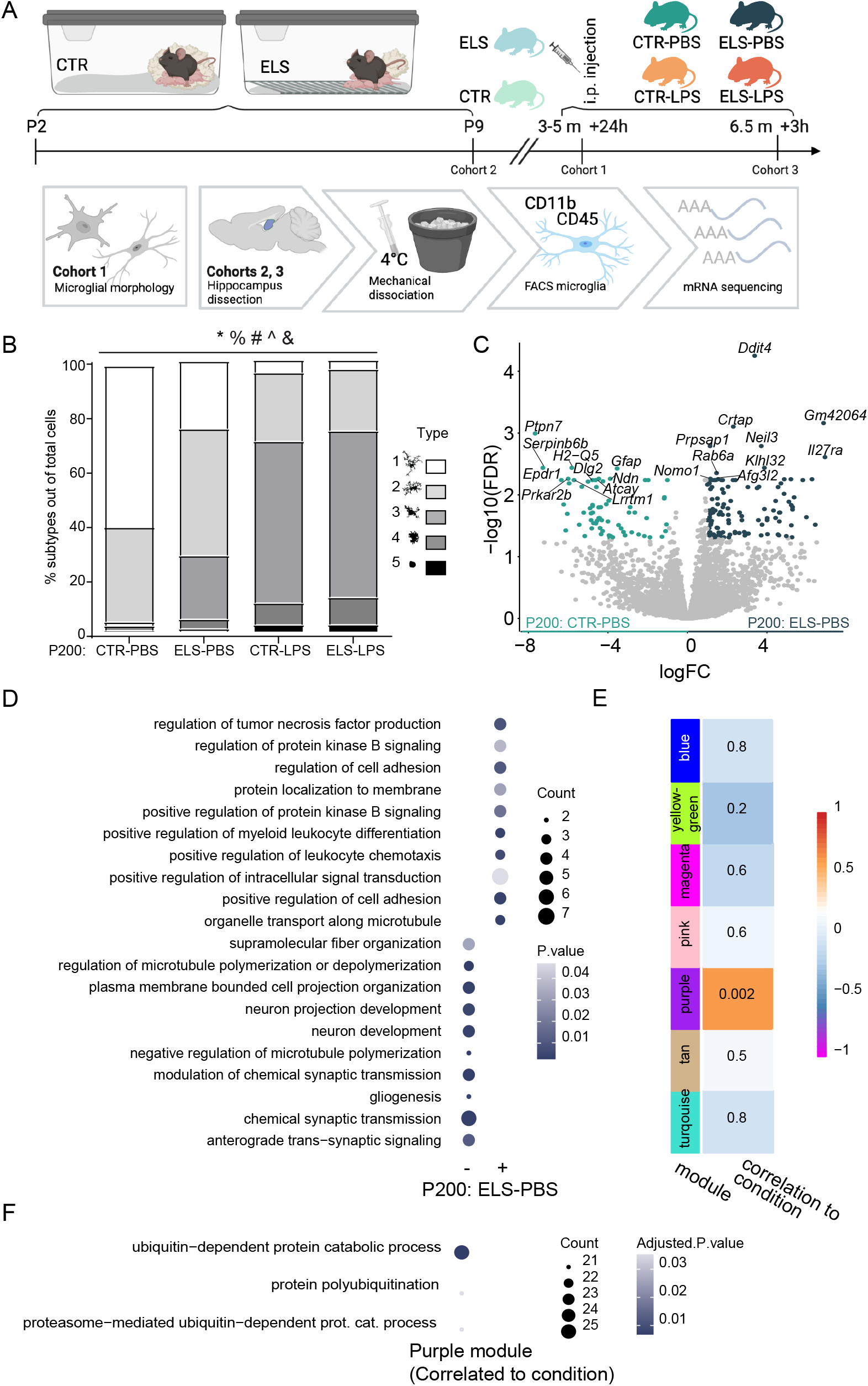
ELS exerts long-term, but no immediate effects on the microglia transcriptome. (A) Overview of the experimental design for the microglia morphometric and transcriptomic analysis (cohorts 1, 2 and 3): from P2 to P9 mice were exposed to limited bedding and nesting material resulting in early-life stress. Control mice were left undisturbed. For the morphological analysis of microglia (cohort 1), mice received an i.p. injection with LPS (5 mg/kg) at 3-5 months (CTR-PBS (n=5), ELS-PBS (n=4), CTR-LPS (n=10), ELS-LPS (n=6) and were sacrificed 24 h later. Two other cohorts were created for transcriptomic analysis of microglia. Cohort 2 was sacrificed immediately after ELS at P9 (12 CTR (n=12), ELS (n=13)) and cohort 3 was left undisturbed until P200, when they received an i.p. injection with PBS or LPS (1 mg/kg) (CTR-PBS (n=7), ELS-PBS (n=6), CTR-LPS (n=7), ELS-LPS (n=5) and were sacrificed 3 hours later. From cohort 2 and 3, hippocampi were dissected, microglia were FACS-purified, and expression profiled. Created with BioRender.com (B) Effects of condition (CTR/ELS) and treatment (PBS/LPS) on the proportion of morphological microglia subtypes in the hilus of 3-5 months mice. General Linear Model Multivariate test, *: main effect treatment, %: interaction effect treatment*condition, #: treatment effect for subtype 1, 3 and 4, ^: condition effect for subtype 1, &: interaction treatment*condition for subtype 1. p<0.05. (C) Volcano plot depicting differentially expressed genes between P200: ELS-PBS and P200: CTR-PBS (logFC</>1, FDR<0.05). Each dot represents a gene. Significantly more abundant genes in P200: CTR-PBS are marked in turquoise and significantly more abundant genes in P200: ELS-PBS are marked in dark turquoise. The 10 most enriched genes in each condition are labelled. (D) Gene ontology (GO) analysis of relatively lower (−) and abundant (+) significant genes in P200: ELS-PBS when compared to P200: CTR-PBS. Top 10 significantly enriched GO terms (p<0.05) based on gene count are depicted (Table S5). (E) Pearson correlation of modules detected with weighted gene co-expression network analysis and condition (ELS/CTR). P-value is indicated as number and R^2^ as color for each correlation. (F) Significantly enriched GO terms associated with the pink module genes (adjusted p-value<0.05, Table S7). Abbreviations: CTR = control, ELS = early-life stress, GO = gene ontology, h = hours, LPS = lipopolysaccharide, m = months, PBS = phosphate-buffered saline.

For a functional readout of microglial function, we extracted microglia from cohorts 4 and 5, which were kept until P9 or P200, respectively (*CTR-P9*: n=8, *ELS-P9*: n=12, *CTR-P200*: n=8, *ELS-P200*: n=3). The extracted microglia were incubated with age-matched synapses isolated from cohorts 6 and 7, which were sacrificed at P9 and P150, respectively (*CTR-P9-synapses*: n=12, *ELS-P9-synapses*: n=11, *CTR-P150-synapses*: n=8, *ELS-P150-synapses*: n=7).

All experimental procedures were conducted according to the Dutch national law and European Union directives on animal experiments and were approved by the Animal Welfare Body of the University of Amsterdam.

### Lipopolysaccharide injection

Adult mice were intraperitoneally (i.p.) injected with either Dulbecco’s Phosphate-Buffered Saline (PBS, Sigma Aldrich D8537) or lipopolysaccharide (LPS, Sigma-Aldrich, *E. coli*, O111:B4, L4391) dissolved in PBS at a dose of 5 mg/kg (cohort 1) or 1 mg/kg (cohort 3) body weight. Twenty-four (cohort 1) or three hours (cohort 3) after injection, blood was collected via tail cuts and mice were sacrificed with i.p. euthasol injection and transcardial perfusion.

### Immunohistochemistry

Microglia were detected in free-floating paraformaldehyde (PFA)-perfused brain tissue by targeting ionized calcium binding adaptor molecule 1 (rabbit anti-IBA1, 019-19741, Wako, see Supplementary Methods for detailed immunohistochemistry protocol).

Morphological analysis was done at 10x magnification on a Nikon Eclipse Ni-E microscope. Cell density was obtained by counting the number of IBA1^+^ cell bodies in the dentate gyrus and cornu ammonis and normalizing to total area, while coverage was measured by dividing the thresholded IBA1 signal by the total area. Microglia in the hilus of the dentate gyrus and the stratum lacunosum-moleculare of the cornu ammonis were classified into five morphological phenotypes as previously described (14,16,41).

Coverage and density data were analyzed using a two-way ANOVA, while the morphological subtypes were analyzed with a general linear multivariate model, all using SPSS 25 (IBM software) and graphed using ggplot2 (v3.3.3.9000)(42) in R. Data were considered statistically significant when p<0.05. Analyses were performed by a researcher blind to animal conditions.

### Microglia isolation and mRNA sequencing

Microglia were isolated from left and right hippocampi of saline-perfused mice as described (43). Briefly, after mechanical dissociation and myelin removal in adult (but not P9) brains using Percoll (Cytiva, 17-5445-02), cells were incubated with a blocker for the mouse Fc Receptor (5 µg/ml, eBioscience, 14-0161) for 15 min. Afterwards, cells were stained with anti-mouse CD11b-PE (1.2 µg/ml, eBioscience, 12-0112) and anti-mouse CD45-FITC (2.5 µg/ml, eBioscience, 11-0451) for 30 min. Shortly before cell sorting, DAPI (0.15 µg/ml, Biolegend, 422801) and DRAQ5 (2 µM, Thermo Scientific, 62251) were added to the cell suspension. Single, viable (DAPI^-^, DRAQ5^+^) microglia (CD45^int^, CD11b^+^) were sorted with the Beckman Coulter MoFlo XDP and were collected in 350 µl RNA lysis buffer (Qiagen, 1053393), and stored at −80°C until further use. Following RNA isolation, mRNA sequencing, and sequence alignment after quality check, bioinformatic analyses were performed in RStudio (v4.0.2). Differential gene expression was determined by a log-fold change of 0.1 and FDR <0.05. Gene ontology analysis was performed using enrichR (v3.0)(44) based on the “GO_Biological_Process_2021” database. Of the cited GO terms distinctly overrepresented in CTR or ELS groups, we also highlighted their associated genes; for terms with >4 associated genes, the top 5 are listed based on absolute logFC (see Supplementary Methods for further details on microglia isolation, mRNA sequencing, and downstream analyses).

### Ex vivo synaptosome phagocytosis assay

We adapted a flow cytometry based ex vivo phagocytosis assay (45). After sacrifice via rapid decapitation, microglia were enriched from whole brains (P9) or half brains from the cortex until the midbrain (P200) using an isotonic Percoll gradient (Supplementary Methods). We incubated 50,000 (P9) or 80,000 (P200) cells with age-matched hippocampal synaptosomes from P9 (1.2 µg) or P150 (2 µg) mice in 300 µl DMEM-F12. We used one tube per mouse as negative control to ensure signal specificity. Synaptosomes were extracted based on a published protocol (46) (Supplementary Methods), and were conjugated to pHrodo-red (P36600, Invitrogen) according to manufacturer instructions. Staining was performed by first blocking the mouse Fc Receptor (5 µg/ml, eBioscience, 14-0161) for 15 min and then by incubating with anti-mouse CD11b-APC (1 µg/ml, eBioscience, 17-0112-82) for 30 min. DAPI (0.15 µg/ml, Biolegend, 422801) was added before flow cytometry analysis using the BD FACS Diva (BD Biosciences).

Approximately 1500 DAPI^-^/CD11b^+^ cells were recorded per tube, and phagocytosis was defined as the proportion of CD11b^+^pHrodo^+^ cells divided by the total number of DAPI^-^/CD11b^+^ cells. Data analysis was done using a mixed linear model using the *nlme* package in R (47), correcting for the seeding of multiple tubes from each animal, as well as nest effects in P9 samples.

### Human cohort and fluorescent *in situ* hybridization in post-mortem human hippocampus

Fresh-frozen hippocampal tissue, from well-characterized age-matched adult males, (depressed suicides with a history of child abuse, n=7, and matched sudden-death controls, n=6) were obtained from the Douglas-Bell Canada Brain Bank (Montreal, Canada). Characterization of early-life histories was based on adapted Childhood Experience of Care and Abuse interviews assessing experiences of sexual and physical abuse (see Supplementary Methods for further details on human cohort). Group characteristics can be found in Table S1, together with correlations between covariates (age, post-mortem interval (PMI), pH, substance dependence, and medication) and the variables measured in this study.

Hippocampal tissues were cut into 10⍰µm-thick sections with a cryostat and collected on Superfrost charged slides. *In situ* hybridization was performed for Hs-TMEM119 and Hs-GAS6 using Advanced Cell Diagnostics RNAscope® probes and reagents following the manufacturer’s instructions (see Supplementary Methods for further details). Sections were imaged using Olympus VS120 virtual slide microscope at 20x magnification. Dentate gyrus area was demarcated manually and QuPath (v0.3.2) (48) was employed for automated cell detection based on DAPI (Vector Laboratories) staining and RNAscope signal quantification. For each probe, cells bearing 3 or more fluorescence puncta were counted as positive.

## Results

### Morphological characterization of hippocampal microglia from ELS-exposed mice under basal and immune-challenged conditions

Microglia can adapt a range of morphologies in response to homeostatic disbalance (16,49,50). We determined the effect of ELS on microglia density, coverage and morphology in subregions of the hippocampus of adult mice under basal conditions and in response to LPS (Figure 1A). ELS exposure did not affect IBA1+ cell numbers or coverage in the dentate gyrus and the cornus ammonis subregions (Figure S1B,C). However, LPS increased IBA1^+^ cell density in both areas (DG: F=27.179, p<0.001; CA: F=23.821, p<0.001,) and reduced IBA1^+^ coverage especially in the dentate gyrus (F=5.528, p=0.028; cornus ammonis: F=4.223, p=0.053, Figure S1D,E).

To further investigate microglial morphology, we characterized microglial subtypes (16,41,51) (Figure S1F) in the hilus and stratum lacunosum-moleculare (SLM). We identified two main morphological subtypes in PBS-injected mice, characterized by either a small cell soma and long branched ramifications (type 1) or a larger cell soma and thicker, branched ramifications (type 2). In PBS-injected animals, ELS decreased the proportion of type 1 microglia in the hilus (interaction treatment*condition: F=9.621, p=0.006; main effect condition: F=11.135, p=0.003) and SLM (main effect condition: F=8.606, p=0.008). Additionally, two other subtypes (type 3 and 4) were observed mostly in ELS mice, characterized by fewer ramifications and larger cell bodies than subtypes 1 and 2. Number of Type 3 microglia was increased by ELS in PBS-injected mice in both regions (Figure 1B, Figure S1G).

LPS significantly affected the proportions of morphological subtypes in the hilus (GLM main effect treatment, F=8.386, p<0.001, Figure 1B) and SLM (GLM main effect treatment, F=8.386, p<0.001, Figure S1G), with modulation by ELS in the hilus (GLM interaction effect treatment*condition, F=3.181, p=0.035, Figure 1B). LPS reduced type 1 microglia in the hilus (F=49.442, p<0.001) and SLM (F=50.609, p<0.001) and increased type 3 (hilus: F=20.789, p<0.001; SLM: F=33.000, p<0.001) and type 4 (hilus: F=8.537, p=0.008; SLM: F=5.719, p=0.027) microglia regardless of early-life condition. A fifth subtype, characterized by an amoeboid morphology, was also detected in some LPS-injected mice (F=3.171, P=0.090) (Figure 1B, Figure S1G).

In summary, under basal conditions ELS altered the proportion of morphological subtypes associated with immune reactivity. Expectedly, LPS treatment also induced a morphological profile associated with inflammation, independent of early-life condition. In this experiment, a relatively high LPS dose was used (5 mg/kg, i.p.), which might have overruled the potentially subtler effects of ELS on microglia. To better detect these modulatory effects of ELS on LPS responses, we used a dose of 1mg/kg LPS for the transcriptomic experiment.

### ELS impacts the microglia transcriptome on the long-term at P200 but not immediately after stress exposure at P9

To determine the acute (P9) and long-term (P200) effects of ELS and LPS on microglial gene expression, we performed mRNA sequencing on purified hippocampal microglia (Figure 1A, Figure S2A). We confirmed the purity of sorted microglia by the high expression of microglial signature genes, but not of other brain cell types (Figure S2B, Table S2). Correlation analysis of the first six principal components (PC) with the experimental variables revealed “age” as the main source of variability in the dataset (with PC1, R^2^=0.97, FDR<0.001), followed by “treatment” (with PC2, R^2^=0.23, FDR<0.01; with PC3, R^2^=0.67, FDR<0.001), and “condition” (with PC6, R^2^=0.15, FDR<0.05; Figure S2C).

When comparing microglial transcriptomes between CTR and ELS-exposed animals at P9, almost no transcriptional changes were found. Only one differentially expressed gene (DEG) was detected, triggering receptor expressed on myeloid cells (*Trem1*) (logFC</>1, FDR<0.05, Figure S2D, Table S3). This gene was however differentially expressed in only 3 out of a total of 13 ELS-exposed mice (Figure S2E) and was therefore not considered biologically relevant for ELS.

In adulthood, we detected 186 DEGs when comparing gene expression profiles of CTR and ELS-exposed animals at basal state, injected with PBS (*P200: ELS-PBS* versus *P200: CTR-PBS)*, (Figure 1C, Table S4). Gene ontology (GO) analysis revealed that the genes downregulated by ELS were involved in “regulation of microtubule (de)polymerization/plasma membrane bounded cell projection organization/neuron development” (*Map, Stmn2, Stmn3*) and “gliogenesis” (*Cdh2, Metrn*). Genes upregulated by ELS were associated with inflammatory pathways and processes, such as “regulation of tumor necrosis factor production” (*Ripk, Gas6, Trim27, Pf4*), “regulation of protein kinase B signaling” (*C1qbp, Gas6, Fermt2, Myorg*), “positive regulation of leukocyte chemotaxis” (*Akirin1, C1qbp, Gas6*), and “positive regulation of cell adhesion” (*Frmd5, C1qbp, Rell2, Dusp26*) (Figure 1D, Table S5).

To identify modules of genes with similar expression patterns in an unbiased manner, Weighted Gene Co-expression Network Analysis (WGCNA) was performed on all P200 samples (*P200: CTR-PBS, P200: ELS-PBS, P200: CTR-LPS, P200: ELS-LPS*). Thirteen co-expression modules were identified (Figure S2F), of which one (purple, Table S6) significantly correlated with early-life condition (R^2^= 0.6, p=0.002, E). GO analysis of genes in this module suggest a role in protein ubiquitination (Figure 1F, Table S7).

In brief, ELS does not impact the microglial transcriptome at P9, but at P200 we detected an upregulation of genes associated with inflammatory processes and protein ubiquitination and a downregulation of genes linked to morphological reconstruction.

### Shared and unique transcriptional changes between P9 and P200 CTR and ELS microglia

To investigate how ELS impacts microglial transcriptional changes over development from P9 to P200, we compared the transcriptomes of P9 and P200 microglia from CTR and ELS-exposed animals, revealing DEGs shared between (grey dots, 2899) and unique for CTR and ELS microglia (*P9: ELS*, light blue dots, 617; *P200: ELS-PBS*, dark turquoise dots, 473; *P9: CTR*, light green dots, 797; *P200: CTR-PBS*, turquoise dots, 353) (Figure 2A, Table S8 and S9). These developmental DEGs (P200 compared to P9) in CTR and ELS animals mostly did not overlap with the transcriptional changes between *CTR-PBS* and *ELS-PBS* animals at P200, pointing to an altered maturation gene expression program of mouse microglia in response to ELS (*Error! Reference source not found*.B).

**Figure 2:**
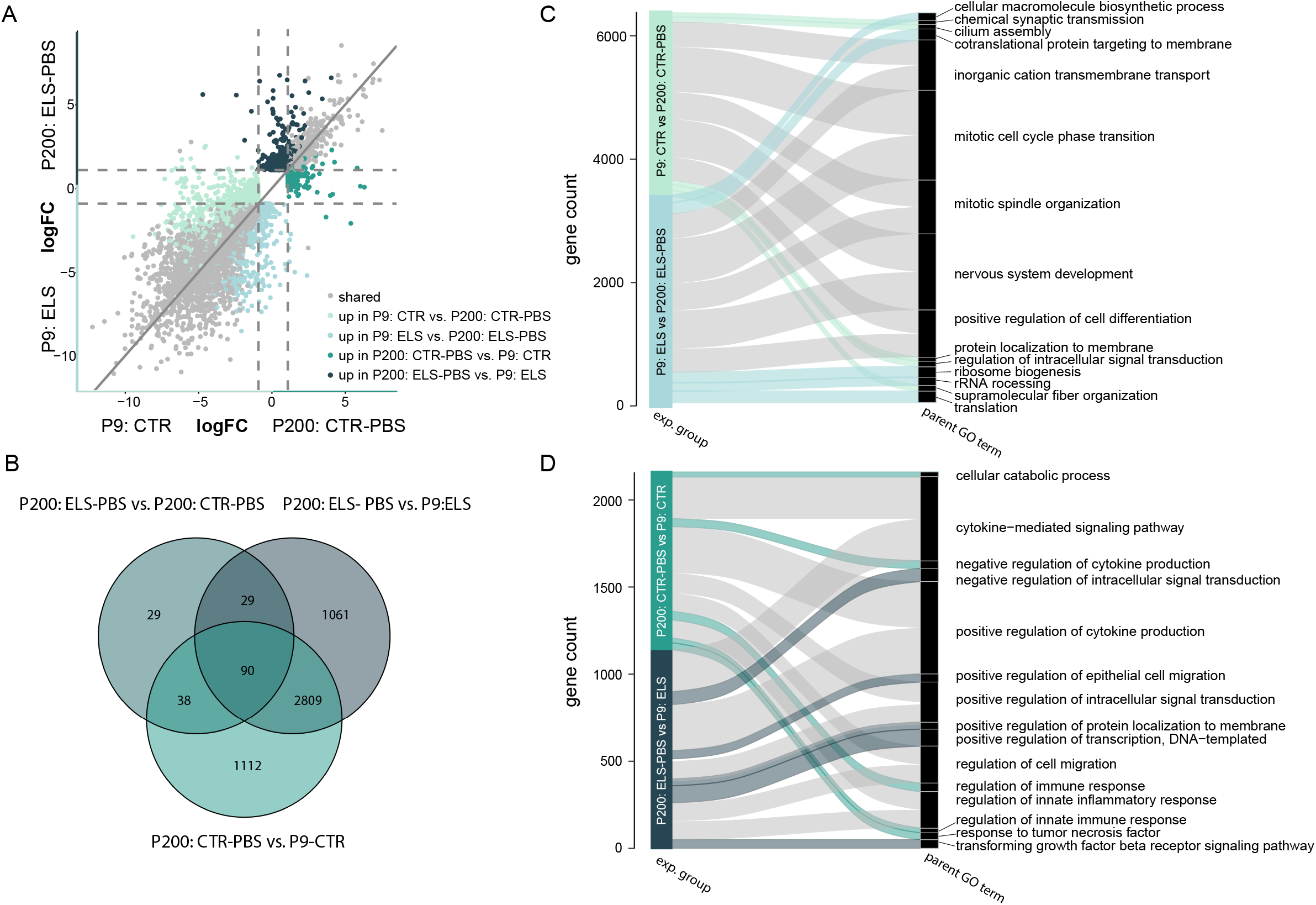
ELS alters microglia development from P9 to P200. (A) Four-way plot depicting expression changes in developmental (P200 vs. P9)-associated genes in ELS and CTR microglia (logFC</>1, FDR<0.05, Table S8 and S9). Each dot represents a gene. Light green dots mark DEGs uniquely enriched in CTR microglia of P9 compared to P200 animals. Light blue dots mark DEGs uniquely enriched in ELS microglia of P9 compared to P200 animals. Turquoise dots mark DEGs uniquely enriched in CTR microglia of P200 compared to P9 animals. Dark turquoise dots mark DEGs uniquely enriched in ELS microglia of P200 compared to P9 animals. Overlapping genes in the development from P9 to P200 of ELS and CTR microglia are marked in grey. (B) Venn diagram depicting the gene expression overlap of all DEGs between adult ELS effects (ELS vs. CTR, Table S3) and developmental effects (P200 vs. P9) in ELS and CTR microglia. (C, D) Alluvial plots illustrating the top 5 enriched parent GO terms P9-(C) and P200-associated (D) genes in CTR and ELS microglia. Significant GO terms (p<0.05) for each experimental group were reduced into parent GO terms (Table S10 and S11), which were ranked based on the total gene count belonging to that parent GO term. Color indicates the experimental group and ribbon thickness depicts the number of genes overlapping with parent GO term-specific genes. Abbreviations: CTR = control, ELS = early-life stress, exp.= experimental, GO = gene ontology, PBS = phosphate-buffered saline.

GO analysis was performed on the shared and unique DEGs and redundant GO terms were reduced into parent GO terms (Figure 2C,D, Table S10 and S11). The shared DEGs in CTR and ELS microglia revealed that P9 microglia had relatively enriched expression of genes associated with mitosis and neurodevelopmental processes (e.g., “mitotic cell cycle phase transition”, “positive regulation of cell differentiation”, “nervous system development”, “chemical synapse organization”; Figure 2C, grey ribbons), whereas at P200, microglia upregulated genes associated with inflammation (e.g. “positive regulation of cytokine production”, “regulation of cell migration”, “regulation of innate inflammatory response”, Figure 2D, grey ribbons).

In CTR animals specifically, P9 microglia were uniquely enriched in genes associated with the “regulation of intracellular signal transduction” (e.g. *Gnai1, Mapk11, Arhgef17, Bst1, Rhov*), “supramolecular fiber organization” (e.g. *Shroom2, Col9a3, Tspan15, Myo5b, Ccdc13*) and “chemical synaptic transmission” (e.g. *Chrna4, Crhbp, Hrh1, Pcdhb16, Pcdhb5*) (Figure 2C, light green ribbons), while P200 microglia were uniquely enriched in genes controlling the immune response (e.g., “response to tumor necrosis factor (TNF)”: e.g. *Acod1, Hyal3, Ccl3, Nfe2l2, Ccl25*; “negative regulation of cytokine production”: e.g. *Acod1, Mefv, Ptpn22, Ppp1r11, Fcgr2b*; “regulation of innate immune response”: e.g. *Acod1, Ptpn22, Trim21, Birc3, Polr3f*; Figure 2D, turquoise ribbons).

In ELS-exposed animals, P9 microglia were uniquely enriched in genes associated with biological processes such as “cellular macromolecule biosynthetic process” (e.g. *Rps16, Rps15a, Rpl39, Rps27a, Rps12*), “cotranslational protein targeting to membrane” (e.g. *Rps16, Rps15a, Rpl39, Rps27a, Rps12*), “ribosome biogenesis” (e.g. *Rps16, Rpl39, Rpl17, Rps24, Rpl9*) and “rRNA processing” (e.g. *Rps16, Rpl39, Rpl17, Rps24, Rpl9*) (Figure 2C, light blue ribbons), whereas genes uniquely enriched at P200 were related to the “negative regulation of intracellular signal transduction” (e.g. *Ddit4, Gper1, Pik3cb, Prkaa2, Per1*), “positive regulation of transcription, DNA-templated” (e.g. *Gper1, Thrb, Ciita, Nr4a1, Zbtb16*), and “transforming growth factor beta (TGFβ) signaling pathway” (e.g. *Gdf9, Src, Smurf1, Zfyve9, Arhgef18*) (Figure 2D, dark turquoise ribbons).

These observations show that independent of early-life condition, P9 microglia are involved in processes related to cell division, cell differentiation and neurodevelopment, while adult microglia perform more inflammation-related. Additionally, ELS at P9 specifically induced processes related to protein translation and biosynthesis of other macromolecules, and at P200 induced genes related to TGFβ signaling.

### ELS impacts the microglia gene expression response to an LPS challenge in adulthood

To determine if ELS alters the transcriptional response of microglia to a systemic immune challenge, we compared microglial transcriptomes of P200 CTR and ELS mice i.p. injected with either PBS or LPS. We identified shared (grey dots, 810), CTR-specific (*CTR-PBS*, turquoise dots, 299; *CTR-LPS*, orange dots, 423), and ELS-specific (E*LS-PBS*, dark turquoise dots, 253; *ELS-LPS*, dark orange dots, 236) genes dysregulated in response to LPS (Figure 3A, Table S12 and S13).

**Figure 3:**
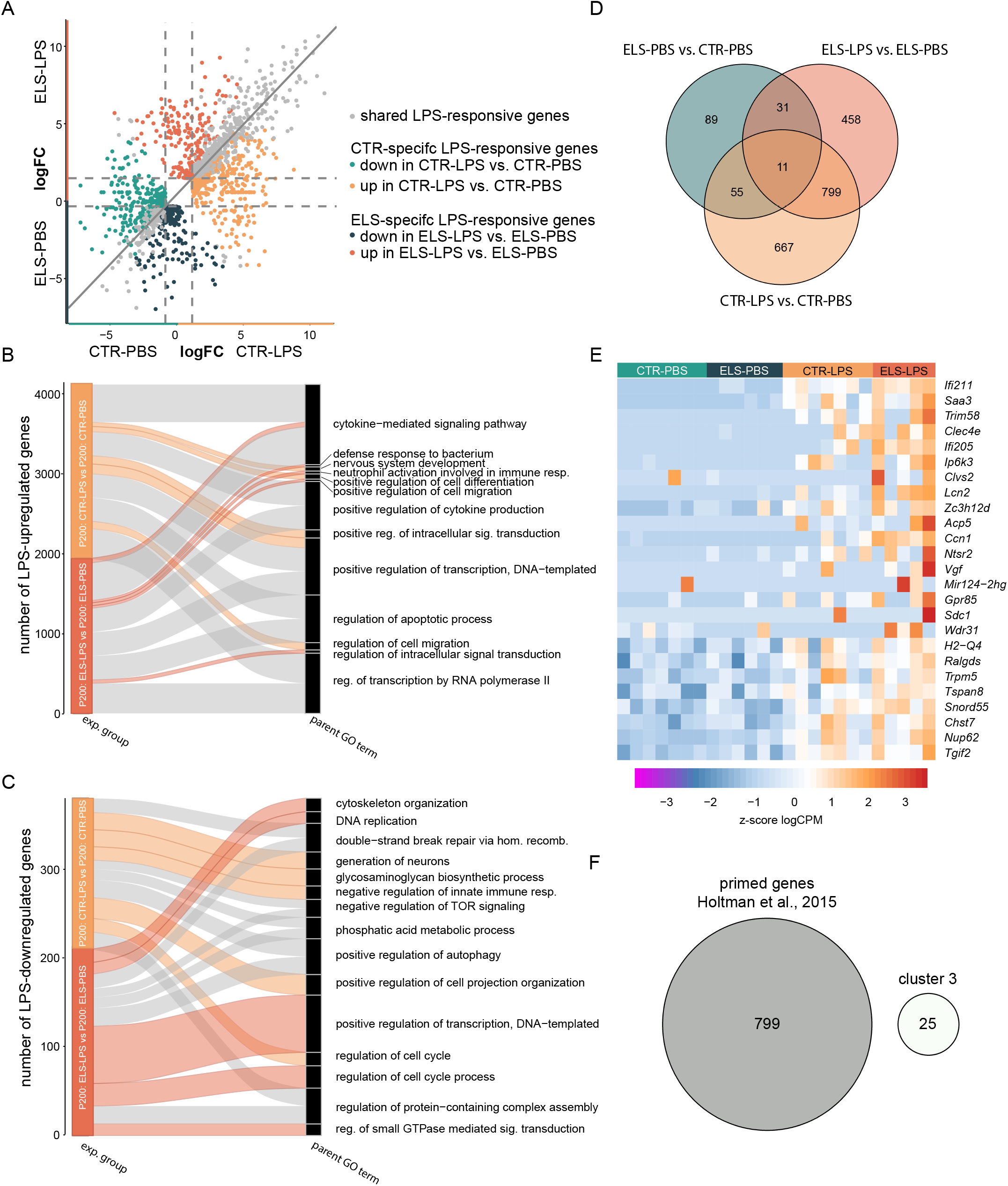
ELS alters the microglia immune response to LPS. (A) Four-way plot depicting gene expression changes in response to LPS (LPS vs. PBS) in ELS and CTR microglia (logFC</>1, FDR<0.05, Tables S12 and S13). Each dot represents a gene. CTR-specific LPS-responsive genes are marked in turquoise and orange, whereby the turquoise dots mark DEGs relatively lowest and orange dots mark DEGs relatively abundant expressed in CTR-LPS compared to CTR-PBS. ELS-specific LPS-responsive genes are marked in dark turquoise and dark orange, whereby the dark turquoise dots mark DEGs relatively lowest and the dark orange dots mark DEGs relatively abundant expressed in CTR-LPS compared to CTR-PBS. Overlapping genes in the LPS response of ELS and CTR microglia are marked in grey. (B, C) Alluvial plots illustrating the top 5 enriched parent GO terms, for upregulated (B) and downregulated (C) genes in LPS-compared to PBS-treated CTR and ELS microglia. Significant GO terms (p<0.05) for each experimental group were reduced into parent GO terms (Tables S14 and S15), which were ranked based on the total gene count belonging to that parent GO term. Color indicates experimental group and ribbon thickness depicts the number of genes overlapping with parent GO term-specific genes. (D) Venn diagram depicting the gene expression overlap between ELS-(P200: ELS-PBS vs. P200: CTR-PBS, Table S3) and LPS-(P200: ELS-LPS vs. P200: ELS-PBS and P200: CTR-LPS vs P200: CTR-PBS) introduced transcriptomic differences in P200 ELS and CTR microglia. (E) Heatmap depicting z-scores of logCPM of primed/trained genes (cluster 3, Figure S3, Table S16). (F) Venn diagram showing gene overlap between genes detected to be primed/trained by ELS and LPS (cluster 3, Table S16) and trained genes detected by Holtman et al. (53) (Table S2). Abbreviations: CTR = control, ELS = early-life stress, LPS = lipopolysaccharide, PBS = phosphate-buffered saline.

LPS-induced upregulated genes in both conditions, as expected, were associated with inflammatory response GO terms, such as “cytokine-mediated signaling pathway”, “positive regulation of cytokine production”, “regulation of apoptotic process” (**Error! Reference source not found**.Figure 3B, grey ribbons, Table S14). Shared downregulated genes in LPS-exposed CTR and ELS mice were related to GO terms such as “double-strand break repair via homologous recombination”, “positive regulation of autophagy”, and “regulation of protein-containing complex assembly” (Figure 3C, grey ribbons, Table S15).

The majority of LPS-induced DEGs in ELS (*ELS-LPS*) compared to CTR (*CTR-LPS*) microglia did not overlap with the gene expression changes induced by ELS itself (*ELS-PBS* vs *CTR-PBS*) (Figure 3D), indicating that the differential response to LPS in ELS microglia were not simply due to the differential expression profile caused by ELS.

Cluster analysis of LPS-responsive genes shared between CTR and ELS microglia identified six gene clusters (Figure S3, Table S16). Genes in cluster 3 are upregulated by LPS in CTR microglia and even more so in ELS microglia (Figure 3E), indicative of microglia training (29,52) by ELS. The list of trained genes in ELS microglia is distinct from a common training gene set detected in (accelerated) aging, and mouse models of Alzheimer’s disease and amyotrophic lateral sclerosis (Table S2, Figure 3F)(53).

Next, GO analysis was performed on the unique transcriptional changes in CTR and ELS microglia in response to LPS, respectively. LPS-induced genes in CTR microglia were related to “positive regulation of cell differentiation” (e.g. *Snai1, Mapk11, Bmpr1b, Zbtb16, Ctnna2*), “positive regulation of intracellular signal transduction” (e.g. *Nedd4, Adra1a, Fn1, Lck, Fermt2*) and “regulation of cell migration” (e.g. *Plet1, Snai1, Ctnna2, Fermt2, Sema3c*) (Figure 3B, orange ribbons), whereas LPS-downregulated genes in CTR microglia were associated with the “glycosaminoglycan biosynthetic process” (*Pxylp1, Ndst2, Ndst1, Hs2st1, B4galt4*), “negative regulation of innate immune response” (*Susd4, Dcst1, Trim21*) and “positive regulation of cell projection organization” (e.g. *Map3k13, Reln, Grip1, Fut9, Ptprd*) (Figure 3C, orange ribbons).

Genes induced in LPS-treated ELS microglia were involved in inflammatory processes such as “defense response to bacterium” (e.g. *Nos2, Chga, Slpi, Isg15, Optn*), “neutrophil activation involved in immune response” (e.g. *Hp, Tnfaip6, Tarm1, Rab37, Sell*) and “positive regulation of cell migration” (e.g. *Sema3e, Pdpn, Edn1, Sod2, Rhoc*) (Figure 3B, dark orange ribbons), whereas LPS-downregulated genes were associated with “cytoskeleton organization” (*Sema6a, Cecr2, Zmym6, Mast1, Arap3*), “DNA replication” (e.g. *Cdc6, Dna2, Chek1, Dbf4, Polg2*) and “regulation of cell cycle process” (*Chek1, Sbf4, Cul9, Zfyve26*) (Figure 3C, dark orange ribbons).

Summarizing, while we observed shared regulation of genes in response to LPS in both adult CTR and ELS microglia, ELS appears to prime a distinct set of LPS-responsive genes in microglia, resulting in a distinct transcriptional response to LPS.

### ELS microglia phagocytes less synaptosomes ex vivo at P200, but not at P9

To further our understanding of the functional consequences of ELS on microglia and to complement our transcriptomic data, we incubated whole brain microglia from P9 and P200 CTR and ELS mice with labeled age-matched hippocampal synaptosomes from CTR and ELS mice.

While at P9 microglia phagocytosis of synaptosomes did not depend on the origin of microglia (Figure 4B, t(4)=0.910, p=0.414), P200 microglia from ELS mice exhibited decreased phagocytiosis of synaptosomes (Figure 4C, microglia condition: t(9)=-2.226, p=0.050). At both ages, ELS synaptosomes were phagocytosed less than CTR synaptosomes, with no interaction between synaptosome source and microglia source (P9 – Figure 4B, synapse condition: t(58)=-5.720, p<0.001; interaction: t(58)=-0.695, p=0.490; P200 - Figure 4C, synapse condition: t(15)=-4.779, p<0.001; interaction: t(15)=0.425, p=0.677).

**Figure 4:**
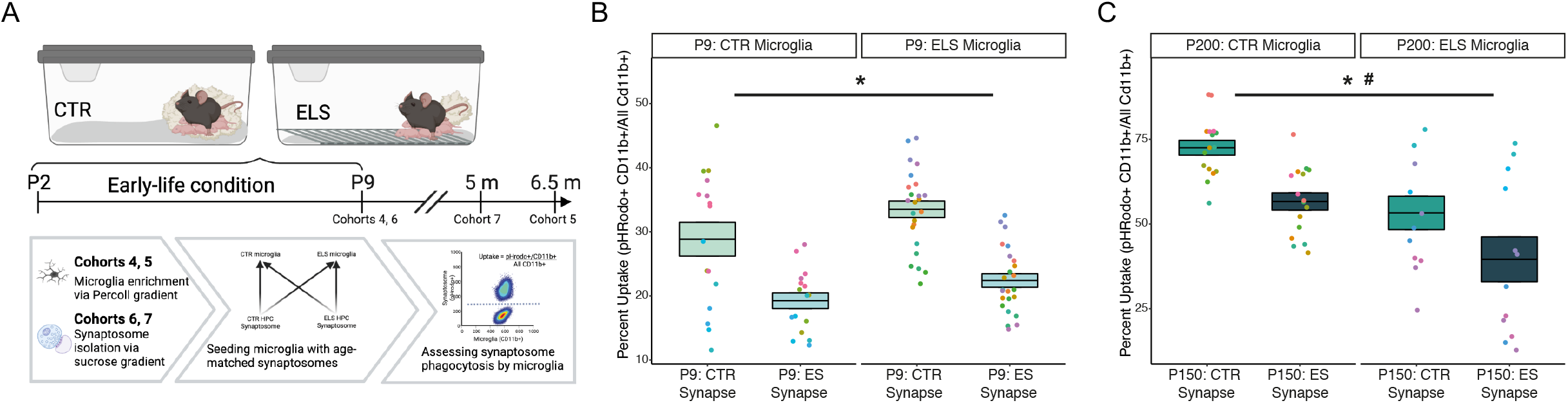
Assessing ELS effects on microglial phagocytosis ex vivo. (A) Experimental design for phagocytosis assay (cohorts 4-7). Whole brain (P9, cohort 4) or hemi-cortex brains (P200, cohort 5) were enriched for microglia using Percoll and incubated with age-matched pHrodo-labelled hippocampal synaptosomes (isolated from cohorts 6-7) for 30 min before flow cytometry analysis. Data were analyzed using a mixed model to correct for animal source and nest for P9 microglia, and animal source for P200 microglia. Created with Biorender.com. (B) Reduced uptake of labelled hippocampal synaptosomes isolated from P9 ELS mice as compared to those from CTR mice, independent of early-life condition of microglia source. (C) At P200, ELS synapses are phagocytosed less than CTR synapses, with a trend towards decreased uptake by ELS microglia. Mixed effect linear model, *: synaptosome-condition effect, p<0.05, #: microglia-condition effect, p=0.05. Abbreviations: CTR = control, ELS = early-life stress, HPC = hippocampus

### Increased microglial GAS6 expression in the hippocampus of post-mortem samples of depressed individuals with a history of childhood abuse

Although animal models are essential to understand the short- and long-term neurobiological consequences of ELS, they cannot fully mirror the complexity of the human brain and experience (54). It is therefore important to study the cellular and molecular consequences of ELS, as experienced through child abuse, in post-mortem human brain samples.

To validate the ELS-induced alterations observed in mice, we selected *Gas6* as a target gene due to its role in phagocytosis (55), its strong upregulation in *ELS-PBS* vs. *CTR-PBS* mice (Figure 5A), and its reported expression also in human microglia (56). We studied *GAS6* in the dentate gyrus (Figure 5B) of post-mortem samples from depressed suicides with a history of childhood abuse (CA, n=7) and matched healthy controls (CTR, n=6) using RNAScope *in situ* hybridization (Figure 5C, D, Supplementary Table). Using a *TMEM119* probe to label microglia, we found that CA subjects displayed increased numbers of microglia (Figure 5E, t(11)=3.308, p=0.007), *GAS6*^*+*^ cells (Figure 5F, t(11)=3.238, p=0.008), and *GAS6*^*+*^ microglia (Figure 5G, t(11)=2.208, p=0.049) in the dentate gyrus. These results highlight the translational value of our ELS mouse model.

**Figure 5:**
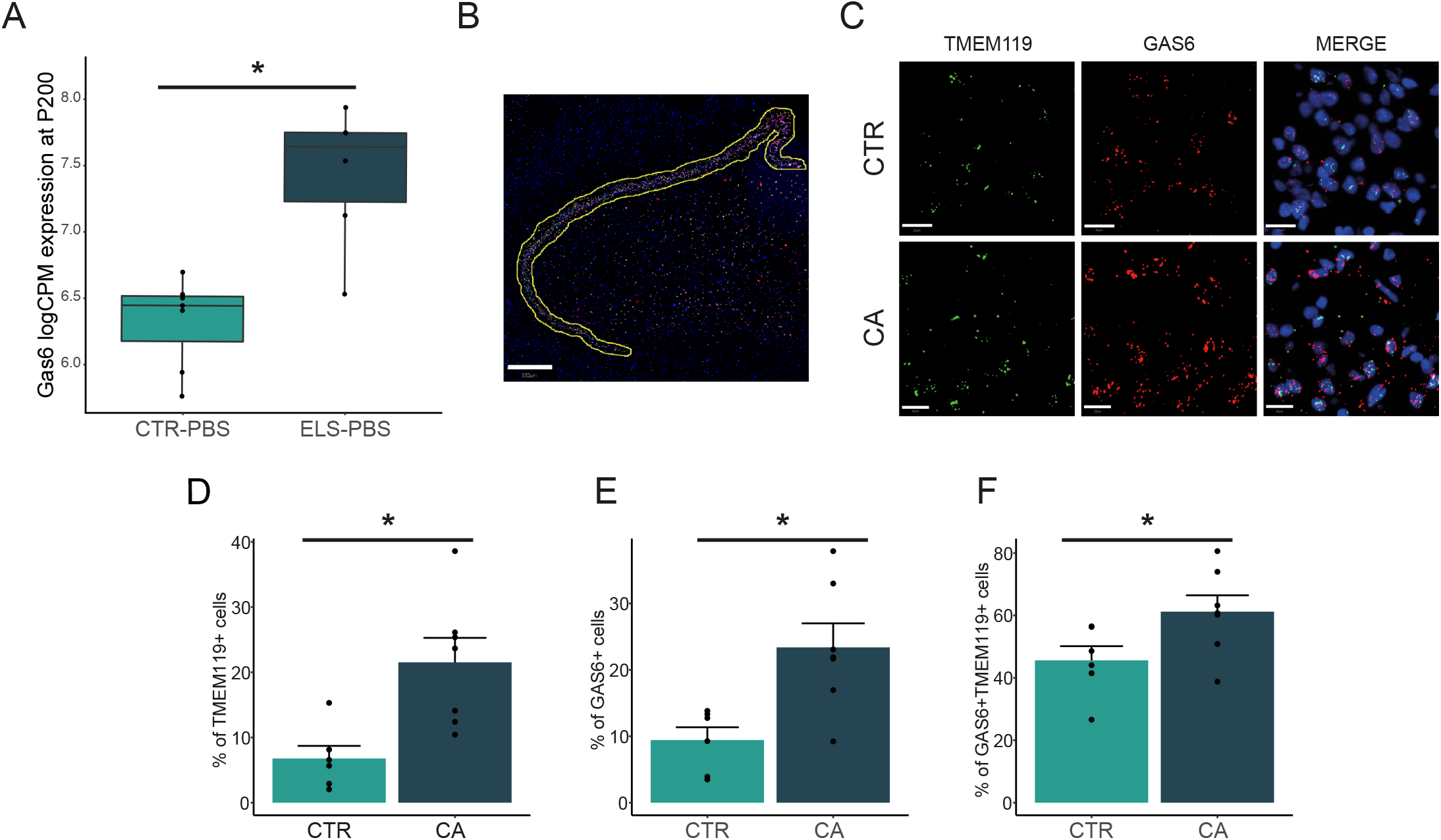
GAS6 expression is increased in mouse and human hippocampal microglia following ELS. (A) Increased expression of microglial Gas6 in mice following early-life stress was validated in (B) human post-mortem dentate gyrus (DG) using RNAscope fluorescent in situ hybridization. Hippocampi from male depressed suicide subjects with a history of childhood abuse (CA, n=7) were compared with age-matched control samples (CTR, n=6). TMEM119 was used to probe microglia. Scale bar: 500µm. (C) Representative image depicting CA-associated changes in the number of cells expressing GAS6, TMEM19 and double positive cells. The number of subpopulations expressing the mRNA of interest were counted and shown as percentage to total cells detected using DAPI staining. Scale bar: 20µm. CA is associated with a significant increase in (D) microglia counts as identified by TMEM19+ cells, (E) total GAS6+ cells, and (F) microglia expressing GAS6 +, detected by cells positive for both TMEM19 and GAS6. Data were analyzed using a t-test. *, condition effect, p<0.05. Abbreviations: CTR: control, CA: childhood abuse.

## Discussion

We demonstrate in this study that ELS exposure in mice, induced by limiting bedding and nesting material, changed the proportion of morphological microglia subtypes and microglial transcriptome in the adult hippocampus, without impacting the transcriptome immediately after stress exposure. Additionally, ELS modulated age-related changes in the microglial gene expression profile between P9 and P200, and the microglia transcriptional response to an LPS challenge in adulthood. The impact of ELS on microglia was also evident at the functional level in the ex vivo phagocytosis of synaptosomes, where P200 (but not P9) ELS microglia exhibited reduced phagocytic capacity. At both ages, ELS synapses were phagocytosed less by both CTR and ELS microglia. Lastly, in order to provide evidence for the translational value of our findings, we demonstrate that one of the identified targets altered by ELS exposure in mouse microglia (i.e. increased GAS6 expression) is also increased in the hippocampal microglia of individuals that experienced childhood adversity.

### ELS affects the microglial transcriptome in adulthood but not immediately after stress exposure

Concerning the immediate effects of ELS on microglia, we did not detect any change in hippocampal microglial transcriptomes after ELS at P9. This is in contrast with the short-term effects on microglial transcriptomes after exposure to brief daily maternal separation (13), as well as with our reported ELS reduction of IBA1 coverage in the dentate gyrus and IBA1+ cell complexity in the hilus at P9 (14). The discrepancy with the findings by Delpech et al. might be due to differences between the ELS model used (brief daily maternal separation versus limited bedding and nesting) and specific ages studied (P14 versus P9). The morphological effects we described previously were subregion-specific (14), while the generated transcriptomic profiles in the present study were from whole hippocampi, possibly diluting out sub-region-specific changes. Alternatively, such transcriptional changes might be latent and emerge only later in life. In fact, there is accumulating evidence for mediation of ELS-associated phenotypes by epigenetic alterations (30,57–59). It is thus possible that ELS-induced effects on microglia at P9 manifest at the epigenetic level, e.g. via stressor- and brain-region-dependent alterations to DNA methylation as described by others (60), that lead to later-life alterations in gene expression and function.

Concerning the long-term effects of ELS on P200 microglia, ELS induced a shift in microglial morphological subtypes, which are indicative for immune reactive cells (51). These alterations were accompanied by increased expression of genes related to inflammatory response, in line with the previously reported ‘immune-activated’ microglial phenotype following ELS (36,38,39), and downregulation of genes involved in microtubule (de)polymerization, typically involved in morphological modulations (61). The motility and dynamics of the microglial cytoskeleton are important for core functions such as chemotaxis (62) and phagocytosis (63), thus potentially contributing to the observed reduction in phagocytic capacity of adult ELS-derived microglia. Furthermore, genes involved in neuronal development, gliogenesis, and microtubule regulation were downregulated by ELS at P200, which could contribute to earlier reported ELS-induced deficits in various forms of cellular plasticity (38,40,64).

With an unbiased WGCNA analysis, we identified an ELS-associated gene module in P200 microglia specifically related to protein ubiquitination and degradation of ubiquitinylated proteins, consistent with reported alterations in the ubiquitin-proteasome system in the hippocampus and cortex of adult rats exposed to maternal separation (65). Ubiquitin is crucial for protein degradation processes and is also involved in pathways such as inflammation (66). This could contribute to ELS-induced increased risk for diseases with a neuroimmune component such as disease, a link that has been proposed in pre-clinical models (67–70) and epidemiological studies (71–73).

Thus, while at P9 there were no apparent effects of ELS on microglial gene expression, these might be latent, emerging later in life to result in the observed long-term programming of microglia impacting the trajectory of gene expression changes required through development and for adaptive response to challenges such as LPS.

### ELS modulates microglia development between P9 and P200

In line with the hypothesis of ELS-associated alterations to developmental trajectories, we observed both shared and unique shifts in gene expression profiles between P9 and P200 in CTR and ELS mice. Microglia upregulated inflammatory pathways across development independent of early-life condition, consistent with earlier reported developmental pattern of microglial gene expression, transitioning from processes related to the cell cycle and pruning to those related to immune surveillance (24,74–77). We detected several differences in the transcriptional changes from P9 to P200 between CTR and ELS animals. For instance, development of CTR microglia specifically involved the Tnf pathway, whereas the Tgfβ pathway was specifically induced during development of ELS microglia. Tgfβ signaling has been reported to drive microglial survival (78), and Tgfβ serum levels have been linked to ELS, as it positively correlated with plasma cortisol levels after an acute stressor in 2 year old primates that experienced ELS (79).

These results indicate that ELS and CTR microglia follow different developmental trajectories, specifically in immune response related genes. The exact implications of this differential trajectory for ELS microglia remain to be determined, but it might underlie how ELS modulates the response to later-life challenges, such as other forms of stress and infection reported by others and us (27,31,32,75,80).

### ELS modulates microglia immune reactivity to LPS challenge in adulthood

Beyond confirming an LPS-induced upregulation of inflammatory genes (22,27,28) and downregulation of autophagy genes (81,82) in both CTR and ELS microglia, we also uncovered a differential LPS response dependent on early-life exposure. These results are in line with the two-hit hypothesis, which postulates that a ‘first hit’ increases sensitivity to later-life challenges, that are unmasked by a ‘second hit’ (83). We previously reported that a first hit in the form of an LPS challenge or deficiency of the DNA repair enzyme Ercc1 interacts with a second LPS hit to unmask long-lasting epigenetic changes leading to either reduced or exaggerated LPS responses as compared to naïve mice injected with LPS (52,84). Here, we demonstrate that ELS can similarly serve as a first hit that affects the transcriptional response of microglia to an LPS challenge in adulthood. Interestingly, beyond exacerbating the expression of a group of LPS responsive genes, suggesting microglia priming (52), ELS also led to a set of distinct transcriptional profile in response to LPS.

### ELS impacts on microglial phagocytosis of synaptosomes

In line with our transcriptomic data, we demonstrated deficits in the ex vivo uptake of synaptosomes in P200, but not P9, microglia isolated from ELS mice. This is important considering the key role of microglial phagocytosis (i.e. pruning) in brain development and function (25,85), and the role of neuron-microglia crosstalk in mediating the hippocampal response to stress (86,87).

At P9, we found reduced microglial phagocytosis, which was driven by the origin of the synaptosomes (i.e. the early-life condition of the mice we extracted synaptosomes from) rather than the origin of the microglia. This suggests that ELS leads to an altered molecular signature in the developmental synapses in the hippocampus. Such reduction of the phagocytosis of ELS synapses, is in line with ELS-induced impaired pruning of excitatory synapses in the hypothalamus at P9 (88). These synaptic signatures could drive the lasting microglial (mal-) adaptations, as we found in our adult data, where the early-life condition of both synaptosomes and microglia contributed to reduced synaptic uptake, possibly ultimately contributing to the well-established ELS effects on hippocampal plasticity (40,89,90). The downregulation of synaptic phagocytosis by P200 ELS microglia might seem counterintuitive considering the literature on ELS-induced reduction of dendritic and synaptic complexity (91–93), which would imply increased phagocytosis. In fact, we observed upregulation of genes regulating phagocytosis (e.g. C1qbp and Gas6) in our adult transcriptomic data. This is in line with the increased phagocytosis of bacterial particles observed in mice exposed to maternal separation (13), even though this was at a much younger age. Our data, taken together with the reported findings by Delpech et al. (13) and Bolton et al. (88), suggest that ELS effects of phagocytosis, rather than being generalized, are more complex, dependent e.g. on specific substrates and possible eat-me signals. What these different pathways are, their regulators, and how ELS modulates them, is still unexplored and awaits future studies.

One specific gene that we explored further is the opsonin *Gas6*. We found that *Gas6* was increased not only in ELS-exposed microglia in mice, but also, importantly, in the post-mortem hippocampi of patients with a history of childhood abuse, both globally and specifically in microglia. GAS6 is a ligand for the tyrosine kinase receptors TYRO3, AXL and MERTK (TAM) that stimulate microglial phagocytosis (94). Its signalling is also known to dampen the LPS inflammatory response of primary cultured microglia (95), mediated by TGF-β expression (96), which was increased in ELS microglia. GAS6 is present at high levels in the brain throughout development, continues to be expressed in adulthood in rodents and humans (56,97), and may act as a neurotrophic factor for hippocampal neurons (98). The modulation of this pathway by ELS is in line with the findings by Bolton et al., where the impaired synaptic phagocytosis of hypothalamic excitatory neurons was mediated via the AXL and MERTK receptors (88). While the increase in GAS6 might raise expectations towards increased microglial phagocytic capacity, it is important to note that GAS6 both activates and is secreted by microglia (55,96). Because activation of the GAS6 receptor MERTK has been shown to stimulate synaptic phagocytosis in astrocytes (99), the observed increase in microglial GAS6 might be a mechanism to recruit other phagocytes to compensate for their deficient functioning.

Moreover, the induction of the TAM pathway by secreted ligands such as GAS6 inhibits prolonged and unrestricted inherent immune responses in macrophages/microglia. Activation of TAM receptors triggers the expression of suppressors of cytokine signalling proteins, which either terminate cytokine receptor-mediated signalling or inhibit nuclear factor kappa B (NF-κB) transcriptional activity (100). In line with this, we found relative downregulation of NF-κB related GO terms in ELS-PBS microglia versus CTR-PBS microglia. According to this mechanism, overexpression of *GAS6* in dentate gyrus microglia after ELS might represent a compensatory mechanism to prevent microglia from becoming hyperresponsive to activation due to stressors or other stimuli in adulthood. This upregulation would imply a non-inflammatory phenotype of microglia in major depression, consistent with recent post-mortem investigations (101) that revealed, e.g., increased homeostatic marker expression in the microglia of depressed individuals (102,103).

An additional significance of the hippocampal GAS6-TAM pathway is its dual role in regulating neurogenesis both directly by supporting neural stem cells, and indirectly by inhibiting microglia and astrocytes (95). Given the ample evidence of long-term modulation of adult hippocampal neurogenesis by ELS (104), there might be clinical implications for targeting TAM receptor-mediated signalling pathways to treat conditions accompanied by neurogenesis loss (100,105), such as depression (101), for which ELS is a major risk factor (3).

Overall, we report here that ELS has long-term effects on hippocampal microglia. ELS altered the distribution of morphological subtypes of microglia, the adult microglia transcriptome at basal state, the microglial developmental trajectory, and their response to an acute immune challenge during adulthood. We provide evidence that these changes have functional consequences for phagocytosis, and that microglia are lastingly impacted in the human brain after childhood abuse. These data highlight the key role of microglia in the lasting effects of ELS exposure, thereby possibly mediating the ELS-induced increased vulnerability to psychopathologies with a neuroinflammatory component such as Alzheimer’s disease and depression.

## Acknowledgement

The authors would like to thank E. Wesseling for assistance during microglia and RNA isolation. The authors would like to acknowledge the expert advice on cell sorting of J. Teunis and G. Mesander from the UMCG Central Flow Cytometry Unit.

## Author contributions

AK, BJLE conceptualized and supervised this study and reviewed and edited the manuscript. KR, LK and JK conceptualized the study and wrote the manuscript. KR and JK performed all mouse-related experimental work. KR and LV performed the morphological characterization of microglia. KR and LK isolated microglia for the sequencing experiments. LK analyzed the sequencing data and prepared the related figures. JK isolated the synaptosomes used for phagocytosis assay, and performed the assay with AvL, and GCS. NM and RR planned the human post-mortem experiments and analyses, which were conducted by RR. GT characterized the post-mortem samples. NB assisted with the isolation of microglia for the sequencing experiments and SMK was instrumental for the analyses and interpretation of RNAseq data. All authors contributed to editing the manuscript.

## Data availability

The mRNA sequencing data generated in this study is currently being upload to the Gene Expression Omnibus database, the link will be made available as soon as possible. All other data will be made available upon request.

## Figure Legends

**Figure S1: ELS validation across cohorts used (related to Materials and Methods: ELS model), and morphological analysis of ELS and control microglia in adulthood under basal and immune-challenged conditions (related to figure 1)**.

(A) ELS decreases body weight gain from postnatal days 2-9 across various cohorts in this project. Body weight at P2 is based on the average body weight, by sex, in each nest. Student’s t-test, *: main effect condition, p<0.05. (B, C, D, E) Iba1+ cell density (B, C) and coverage (D, E) in the dentate gyrus (B, D) and cornu ammonis (C, E) of the hippocampus. Two-way ANOVA, *: main effect LPS, p<0.05; $: trend for LPS effect, p=0.053. (F) Representative images of morphological Iba1+ subtypes. Objective 40x, scale bar = 10 µm. (G) Effects of condition (CTR/ELS) and treatment (PBS/LPS) on the proportion of morphological microglia subtypes in the stratum lacunosum-moleculare of 3-5 months old CTR and ELS-exposed mice. General Linear Model Multivariate test, *: main effect treatment, #: treatment effect for subtype 1, 3 and 4, ^: condition effect for subtype 1. p<0.05. Abbreviations: CTR= control, ELS= early-life stress, ANOVA = analysis of variance, LPS = lipopolysaccharide, PBS = phosphate-buffered saline.

**Figure S2: Transcriptional analysis of ELS and control microglia in adulthood under basal and immune-challenged conditions (related to figure 1)**.

(A) FACS strategy to obtain single, viable (DAPI-, DRAQ5+), CD45+, CD11b+ microglia. (B) Average expression (logCPM) of gene sets specific for different brain cell types (106) (Table S2) in microglia of all experimental groups. (C) Pearson correlation (R2) of the first 6 principal components (PC) to the experimental variables (treatment (PBS/LPS), condition (CTR/ELS) and age (P9/P200)) (FDR ***<0.001, **<0.01, *<0.05). (D) Volcano plot depicting differential expressed genes between P9: ELS and P9: CTR (logFC</>1, FDR<0.05). Each dot represents a gene. Significantly differential expressed gene is labelled and marked with cyan. (E) Gene expression (CPM) of Trem1 in P9: ELS and P9: CTR microglia from individual mice. (F) Gene dendrogram and module colors of weighted gene co-expression network analysis. Abbreviations: AST = astrocytes, CTR = control, CPM = counts per million, ELS = early-life stress, END= endothelial cells, FC = fold change, FSC = forward scatter, LPS = lipopolysaccharide, MIC = microglia, NEU = neurons, OLI = oligodendrocytes, P = postnatal day, PC = principal component, PBS = phosphate-buffered saline, SSC = side scatter.

**Figure S3: Cluster analysis of LPS-responsive genes in CTR and ELS microglia (related to figure 3)**.

Heatmap with Manhattan distance-based hierarchical clustering analysis depicting z-scores of average logCPM values of all genes overlapping between P200: CTR-LPS vs P200: CTR-PBS and P200: ELS-LPS vs P200: ELS-PBS. Abbreviations: CTR= control, ELS= early-life stress, LPS= lipopolysaccharide, PBS= phosphate-buffered saline.

## Supplementary tables

- *Table S1. Clinical information from individuals used in the post-mortem study (related to Materials and Methods: Human cohort and Figure 5).*
- *Table S2. publicly available gene lists of CNS cell types (106) and trained genes (53) doi: 10.1186/s40478-015-0203-5) (related to Figure S2B and Fig 3F).*
- *Table S3. Differential gene expression analysis P9: ELS vs. P9: CTR (logFC</>1, FDR<0.05) (related to Figure S2D,E).*
- *Table S4. Differential gene expression analysis P200: ELS-PBS vs. P200: CTR-PBS (logFC</>1, FDR<0.05) (related to Figure 1C).*
- *Table S5. Gene ontology analysis of genes identified in P200: ELS-PBS vs. P200: CTR-PBS microglia (p value<0.05) (related to Figure 1D).*
- *Table S6. Genes belonging to the pink module identified in WGCNA (related to Figure 1E, Figure S2F).*
- *Table S7. Gene ontology analysis of pink module genes (adjusted p value<0.05) (related to Figure 1F)*
- *Table S8. Differential gene expression analysis P200: CTR-PBS vs. P9: CTR (logFC</>1, FDR<0.05) (related to Figure 2A).*
- *Table S9. Differential gene expression analysis P200: ELS-PBS vs. P9: ELS (logFC</>1, FDR<0.05) (related to Figure 2A).*
- *Table S10. Original (enrichR) and reduced (rrvgo) gene ontology analysis of genes enriched in ELS and CTR microglia in the P200 vs P9 comparison (related to Figure 2C).*
- *Table S11. Original (enrichR) and reduced (rrvgo) gene ontology analysis of genes depleted in ELS and CTR microglia in the P200 vs P9 comparison (related to Figure 2D).*
- *Table S12. Differential gene expression analysis P200: CTR-LPS vs. P200: CTR-PBS (logFC</>1, FDR<0.05) (related to Figure 3A).*
- *Table S13. Differential gene expression analysis P200: ELS-LPS vs. P200: ELS-PBS (logFC</>1, FDR<0.05) (related to Figure 3A).*
- *Table S14. Original (enrichR) and reduced (rrvgo) gene ontology analysis of genes enriched in ELS and CTR microglia in the LPS vs PBS comparison (related to Figure 3B).*
- *Table S15. Original (enrichR) and reduced (rrvgo) gene ontology analysis of genes depleted in ELS and CTR microglia in the LPS vs PBS comparison (related to Figure 3C).*
- *Table S16. Clustering analysis of overlapping genes between the LPS response of ELS and CTR microglia (related to figure 3F).*

