## Supplementary figures and images for "Early-life stress lastingly impacts microglial transcriptome and function under basal and immune-challenged conditions"

### Supplemental figure 1

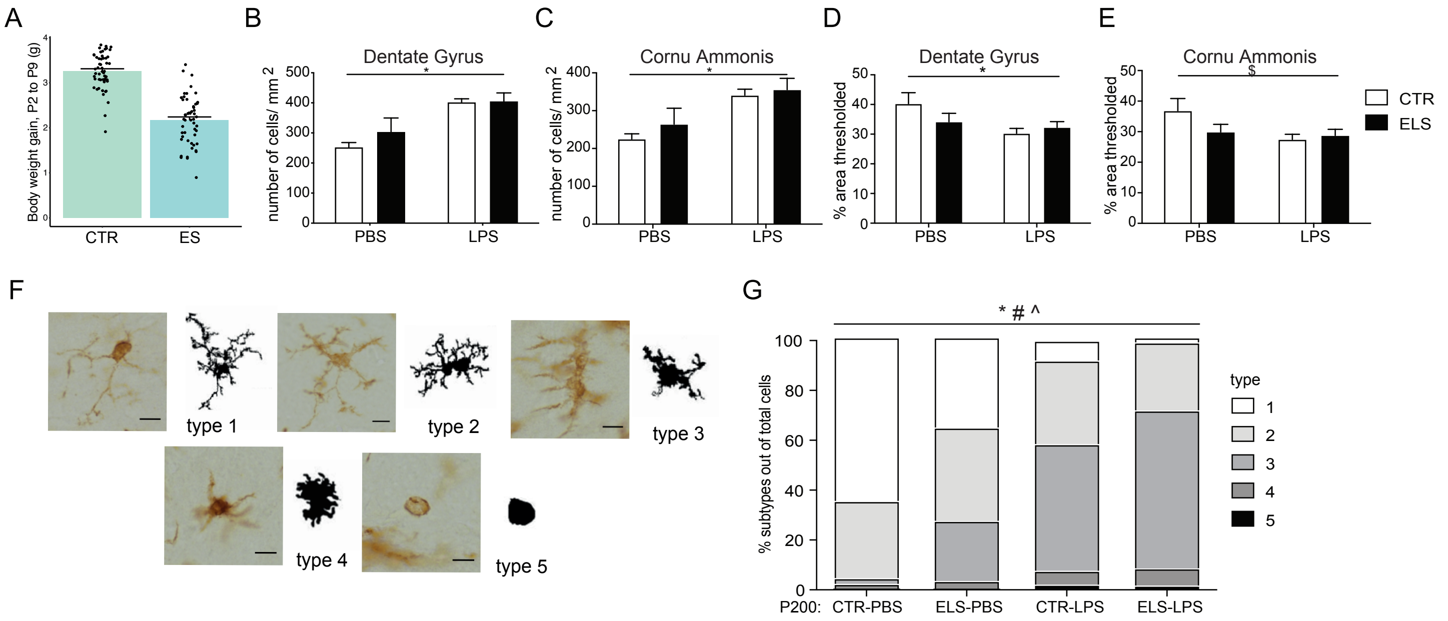

### Supplemental figure 2

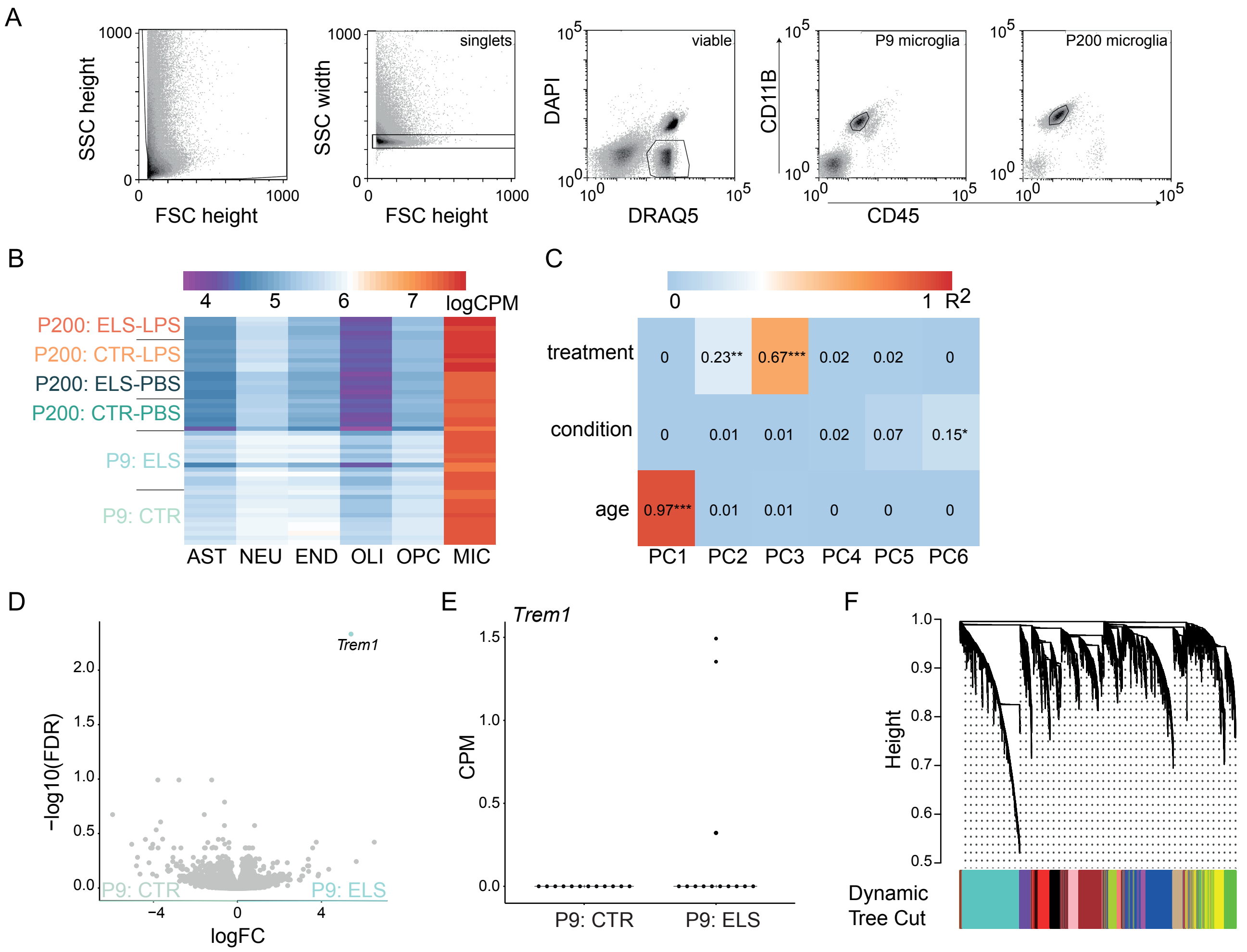

### Supplemental figure 3

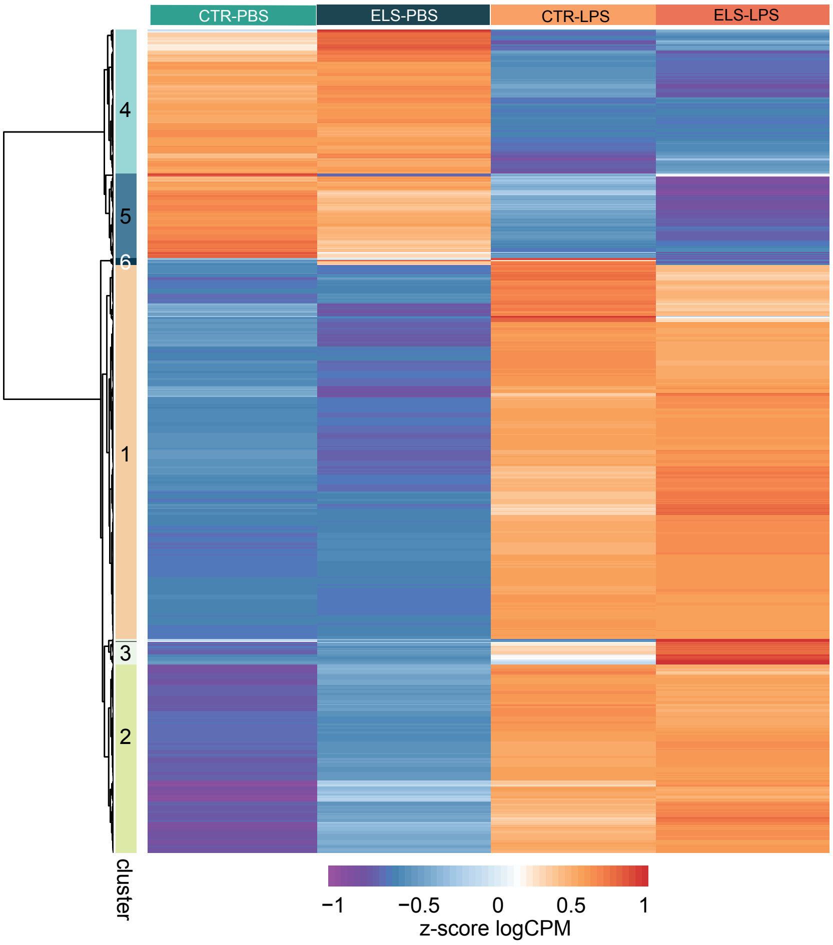
